# 5-HT_2B_ serotonin receptor agonist BW723C86 shapes the macrophage gene profile via AhR and impairs monocyte-to-osteoclast differentiation

**DOI:** 10.1101/587709

**Authors:** Concha Nieto, Ignacio Rayo, Mateo de las Casas-Engel, Elena Izquierdo, Bárbara Alonso, Miguel A. Vega, Ángel L. Corbí

## Abstract

Peripheral serotonin (5-HT) exacerbates or limits inflammatory pathologies through interaction with seven types of 5-HT receptors (5-HT_1-7_). As central regulators of inflammation, macrophages are critical targets of 5-HT, which promotes their anti-inflammatory and pro-fibrotic polarization primarily via the 5-HT_7_-Protein Kinase A (PKA) axis. However, anti-inflammatory human macrophages are also characterized by the expression of 5-HT_2B_, an off-target of anesthetics, anti-parkinsonian drugs and Selective Serotonin Reuptake Inhibitors (SSRI) that contributes to 5-HT-mediated pathologies. Since 5-HT_2B_ prevents mononuclear phagocyte degeneration in amyotrophic lateral sclerosis and modulates motility of murine microglial processes, we sought to determine the functional and transcriptional consequences of 5-HT_2B_ activation in human macrophages. Ligation of 5-HT_2B_ by the 5-HT_2B_-specific agonist BW723C86, which exhibits antidepressant- and anxiolytic-like effects in animal models, significantly modified the cytokine profile and the transcriptional signature in macrophages. Importantly, 5-HT_2B_ agonist-induced transcriptional changes were partly mediated through activation of the Aryl hydrocarbon Receptor (AhR), a ligand-dependent transcription factor that regulates immune responses and the biological responses to xenobiotics. Besides, BW723C86 triggered transcriptional effects that could not be abrogated by 5-HT_2B_ antagonists and impaired monocyte-to-osteoclast differentiation by affecting the expression of negative (*IRF8*) and positive (*PRDM1*) regulators of osteoclastogenesis. Therefore, our results demonstrate the existence of a functional 5-HT_2B_-AhR axis in human macrophages and indicate that the commonly used 5-HT_2B_ agonist BW723C86 exhibits 5-HT_2B_-independent effects. The 5-HT_2B_-AhR link extends the range of signaling pathways initiated upon 5-HT receptor engagement and identifies a point of convergence for endogenous and exogenous agents with ability to modulate inflammatory responses.

**KEY POINTS:**

- The serotonin receptor 5-HT_2B_ modifies the human macrophage transcriptome through activation of the Aryl Hydrocarbon Receptor.
- BW723C86, an agonist used for 5-HT_2B_ activation *in vivo*, exerts 5-HT_2B_-independent effects and limits monocyte osteoclastogenic potential.

## INTRODUCTION

Peripheral serotonin (5-hydroxytryptamine, 5-HT) influences the physiopathology of numerous tissues ^1-3^ as it controls vascular, heart and gastrointestinal functions ^4,5^, promotes cell proliferation ^6-9^, regulates wound healing, and influences immune and inflammatory responses ^10^ by modulating T lymphocyte ^11^ and myeloid cell functions ^12-16^. In inflammatory pathologies, 5-HT contributes to Pulmonary Arterial Hypertension (PAH) ^17^, atopic dermatitis ^18^, systemic sclerosis ^19^, inflammatory gut disorders ^20-25^, cancer angiogenesis ^26^ and neuroendocrine neoplasms proliferation ^27^, whereas it reduces pathologic scores in collagen-induced arthritis ^28^. The existence of seven types of 5-HT receptors (5-HT_1-7_), with different distribution and signaling properties ^29^, underlies these tissue-specific actions of 5-HT.

Macrophages promote the initiation and the resolution of inflammatory processes, with both processes being essential for maintaining tissue homeostasis ^30-32^. In fact, modulation of macrophage polarization is an attractive therapeutic target for chronic inflammatory pathologies, where the balance between pro-inflammatory and resolving macrophages is altered ^33,34^. Not surprisingly, macrophage effector functions are directly modulated by 5-HT ^17,22,26,35,36^, and also by Selective Serotonin Reuptake Inhibitors (SSRI) ^37,38^. We have previously demonstrated that 5-HT_7_ is the major mediator of the anti-inflammatory actions of 5-HT on human macrophages ^39^ and that both 5-HT_2B_ and 5-HT_7_ contribute to the maintenance of the anti-inflammatory state of M-CSF-dependent human macrophages ^40^.

The 5-HT_2B_ serotonin receptor is expressed in the central nervous system, where it exerts anti-depressant or anxiolytic-like effects ^41^, and in periphery, where it plays a major role in heart development ^42^. Pharmacologic and genetic studies indicate that 5-HT_2B_ is required for 5-HT-mediated pathologies like PAH ^17,43^, dermal, lung and liver fibrosis ^19,44,45^, cardiac hypertrophy ^46,47^, and mediates the valvular heart disease (VHD) and fibrosis associated with carcinoid syndrome ^48^. Importantly, anti-migraine drugs like methysergide and ergotamine ^49,50^, general anesthetics ^51^, fenfluramine and conventional SSRIs ^52-55^, and even the dopamine agonists and anti-parkinsonian drugs pergolide and cabergoline ^50,56,57^, exert potent off-target effects on 5-HT_2B_ ^50^. In fact, 5-HT_2B_ appears to be required for the therapeutic actions of SSRI ^54^. Since some of these 5-HT_2B_-targeting drugs induce VHD, their use is restricted, and novel potential drugs are commonly screened for 5-HT_2B_ agonist activity.

5-HT_2B_ engagement enhances proliferation of numerous cell types ^9,58-67^ via Gαq and Src phosphorylation, and through production of growth factors like insulin ^68^, TGFβ1, CTGF, FGF2 ^45,60^ and TGFα ^69^. In macrophages, 5-HT_2B_ prevents mononuclear phagocyte degeneration in amyotrophic lateral sclerosis ^70^, and *Htr2b*^−/−^ microglia exhibits a mild inflammatory state ^71^, which is in line with the ability of 5-HT_2B_ antagonist SB204741 to impair the acquisition of human macrophage polarization-specific genes ^40^ and TGFβ1 expression by mouse Kupffer cells ^45^. However, although 5-HT_2B_ activation modifies inflammatory cytokine production by human monocytes ^72^, the functional and transcriptional consequences of 5-HT_2B_ ligation in human macrophage remains to be determined. Since anti-inflammatory M-CSF-dependent macrophages express 5-HT_2B_, we have assessed the specific role of 5-HT_2B_ in macrophage polarization. The use of the 5-HT_2B_ agonist BW723C86, used to evaluate the role of 5-HT_2B_ in preclinical models of depression ^54^ and anxiety ^73,74^, allowed us to demonstrate that 5-HT_2B_ shapes the macrophage transcriptome via Aryl hydrocarbon Receptor (AhR) activation and impairs the osteoclastogenic potential of human macrophages.

## METHODS

### Generation of human monocyte-derived macrophages and cell culture

Buffy coats from anonymous healthy blood donors were provided by Comunidad de Madrid blood Bank. Human peripheral blood mononuclear cells (PBMC) were isolated from buffy coats over a Lymphoprep (Nycomed Pharma) gradient, and monocytes were purified by magnetic cell sorting using CD14 microbeads (Miltenyi Biotech). Monocytes (>95% CD14^+^ cells) were cultured at 0.5 × 10^6^ cells/ml for 7 days in RPMI supplemented with 10% fetal calf serum (FCS, Biowest) (completed medium) at 37°C in a humidified atmosphere with 5% CO_2_, and containing M-CSF (10 ng/ml) (ImmunoTools GmbH) to generate M-MØ monocyte-derived macrophages. M-CSF was added every two days. Before treatments, M-MØ were maintained in serum-free medium (Macrophage-SFM, Gibco) for 48 hours. For macrophage activation, cells were exposed to 5-HT_2B_ agonists for 6 hours and then treated with *Escherichia coli* 055:B5 LPS (10 ng/ml) for 18 h. The 5-HT_2B_ agonists BW723C86 (α-methyl-5-(2-thienylmethoxy)-1*H*-indole-3-ethanamine) ^75,76^ and α-Methylserotonin (AMS) ^77^, and the 5-HT_2B_ antagonist SB204741 ^78^, were purchased from Sigma-Aldrich. The 5-HT_2B_ agonist 6-APB (6-(2-aminopropyl)benzofuran) was obtained from Cayman and used at the indicated concentrations. The AhR agonist FICZ (Enzo) and antagonist CH223191 (Calbiochem) were used at 250 nM and 3 µM, respectively. When indicated, SB204741 was used 1 hour before treatment with agonists. Monocyte-derived osteoclasts were generated by culturing monocytes for 12 days on glass coverslips with M-CSF (25 ng/ml) and Receptor Activator for NF-kB Ligand (RANKL; 30 ng/ml) addition every 72 h and BW723C86 treatment 6 h before cytokine addition. Osteoclast generation was verified directly by phase contrast microscopy or after staining for Tartrate-Resistant Acid Phosphatase (TRAP) (Leukocyte Acid Phosphatase kit, Sigma-Aldrich). Human TNFα and CCL2 in M-MØ culture supernatants were measured using commercially available ELISA (BD Biosciences).

#### Transfections and Reporter gene assays

Dual luciferase Cignal® Xenobiotic Response Element (XRE) Reporter assay kit (Qiagen), where the XRE-Luc construct harbours tandem repeats of the AhR-binding element (XRE), was used for the reporter gene assays. HepG2 cells (80.000 cells/well) or M-MØ (1×10^6^ cells/well) were transfected with 1 μg of the XRE reporter construct using either Superfect (Qiagen) or VIROMER RED (Lipocalix), respectively. For normalizing transfection efficiency, transfected DNA included a 40:1 mixture of the XRE-specific firefly luciferase construct and a construct expressing *Renilla* luciferase from a constitutive promoter. After transfection, cells were cultured overnight before exposure to BW723C86 (10 μM), FICZ (250 nM), CH223191 (3 μM) or DMSO for 24 h, lysed, and firefly and *Renilla* luciferase activities were determined using the Dual-Luciferase® Reporter Assay System (Promega).

#### Small Interfering Ribonucleic Acid (siRNA) Transfection

To knockdown AhR expression, M-MØ (1 × 10^6^ cells) were transfected with AhR-specific siRNA (siAhR) (50 nM) (# s1199, Thermo-Fisher Scientific), using HiPerFect (Qiagen) using siC (#4390843, Thermo-Fisher Scientific) as negative control siRNA. Cells were allowed to recover from transfection in complete medium (18–24 h) and then treated for 6h with BW723C86 (10 μM) before analysis.

### Quantitative real-time RT-PCR (qRT-PCR)

Total RNA was extracted using the total RNA and protein isolation kit (Macherey-Nagel). RNA samples were retrotranscribed (High-Capacity cDNA Reverse Transcription kit, AB), and triplicates of amplified cDNA were analyzed on a Light Cycler® 480. Gene-specific oligonucleotides (Supplementary Table I) were designed using the Universal ProbeLibrary software (Roche Diagnostics). Results were expressed relative to the expression level of the endogenous reference gene *TBP* and using the ΔΔCT (cycle threshold) method for quantitation.

### Microarray analysis

Global gene expression analysis was performed on RNA from three independent samples of untreated (control), BW723C86-treated (10 μM), SB204741-treated (1 μM), or SB204741+BW723C86-treated M-MØ. RNA was isolated using the RNeasy Mini kit (Qiagen) and used as hybridization probe on Whole Human Genome Microarrays (Agilent Technologies, Palo Alto, CA). Only probes with signal values >60% quantile in at least one condition were considered for the differential gene expression (DGE) and statistical analysis. Statistical analysis for DGE was carried out using empirical Bayes moderated paired t-test implemented in the *limma* package (http://www.bioconductor.org), and p values were adjusted for multiple hypotheses testing using the Benjamini-Hochberg method to control the false discovery rate ^79^, with all procedures coded in R (http://www.r-project.org). Microarray data were deposited in the Gene Expression Omnibus (http://www.ncbi.nlm.nih.gov/geo/) under accession no. GSE121825. Unsupervised hierarchical clustering was done on the mean expression level of genes significantly regulated by BW723C86 (adj *p*<0.002) with the ClustVis webtool (https://biit.cs.ut.ee/clustvis/) ^80^. Differentially expressed genes were analysed for annotated gene sets enrichment using ENRICHR (http://amp.pharm.mssm.edu/Enrichr/) ^81,82^, and enrichment terms considered significant with a Benjamini-Hochberg-adjusted *p* value <0.05. For gene set enrichment analysis (GSEA) ^83^, the gene sets available at the website, as well as the previously defined “Pro-inflammatory gene set” and “Anti-inflammatory gene set” ^84^ (GSE68061) were used.

### Statistical analysis

For comparison of means, and unless otherwise indicated, statistical significance of the generated data was evaluated using the Student paired t-test. In all cases, *p*<0.05 was considered as statistically significant.

## RESULTS

### The BW723C86 agonist modulates the transcriptional signature of human macrophages partially via 5-HT_2B_

Since human anti-inflammatory M-MØ express 5-HT_7_ and 5-HT_2B_ serotonin receptors ^40^ and clinically relevant drugs (anesthetics, SSRIs, antimigraine drugs) act via 5-HT_2B_^49,51-54^, we sought to determine the consequences of 5-HT_2B_ engagement in human macrophages using BW723C86, an 5-HT_2B_ agonist with 10-fold and 100-fold selectivity over human 5-HT_2C_ and 5-HT_2A_, respectively ^85^, amply used as *in vivo* ^54,73,74,86-91^ and *ex vivo* ^28,66,70^, and with potential therapeutic effects ^73,74,91^. Transcriptional profiling of BW723C86-treated M-MØ (Figure 1A) revealed that the agonist increased the expression of 357 annotated genes and downregulated the expression of 398 genes (*adj p<0.002*) (Figure 1B, Supplementary Tables II and III). Analysis of independent M-MØ samples confirmed the microarray data, as BW723C86 increased *NCF4* and reduced *ACSL1*, *EGR1* and *DUSP6* expression (Figure 1C). Gene Set Enrichment Analysis (GSEA) revealed that BW723C86 significantly modifies the expression of “Gα_12/13_ signaling” ^92^ and “Amyotrophic Lateral Sclerosis” gene sets, in agreement with the signaling capability and pathological significance of 5-HT_2B_ ^70^ (Figure 1D), and affects biological processes like “GO Heart Valve Development” and “GO Regulation of Myeloid Leukocyte Differentiation” (Supplementary Figure 1A), in line with the known involvement of 5-HT_2B_ in heart development ^42,93,94^ and macrophage differentiation ^40^. Further, terms associated with bone metabolism were also enriched in BW723C86-treated M-MØ (Supplementary Figure 1A). Interestingly, and unlike the 5-HT-mediated activation of 5-HT_7_ ^39^, BW723C86 significantly augmented the expression of genes associated to both pro-inflammatory and anti-inflammatory macrophage polarization ^84,95^ (Supplementary Figure 1B), suggesting that engagement of 5-HT_2B_ or 5-HT_7_ on human macrophages leads to different transcriptional outcomes.

**Figure 1.**
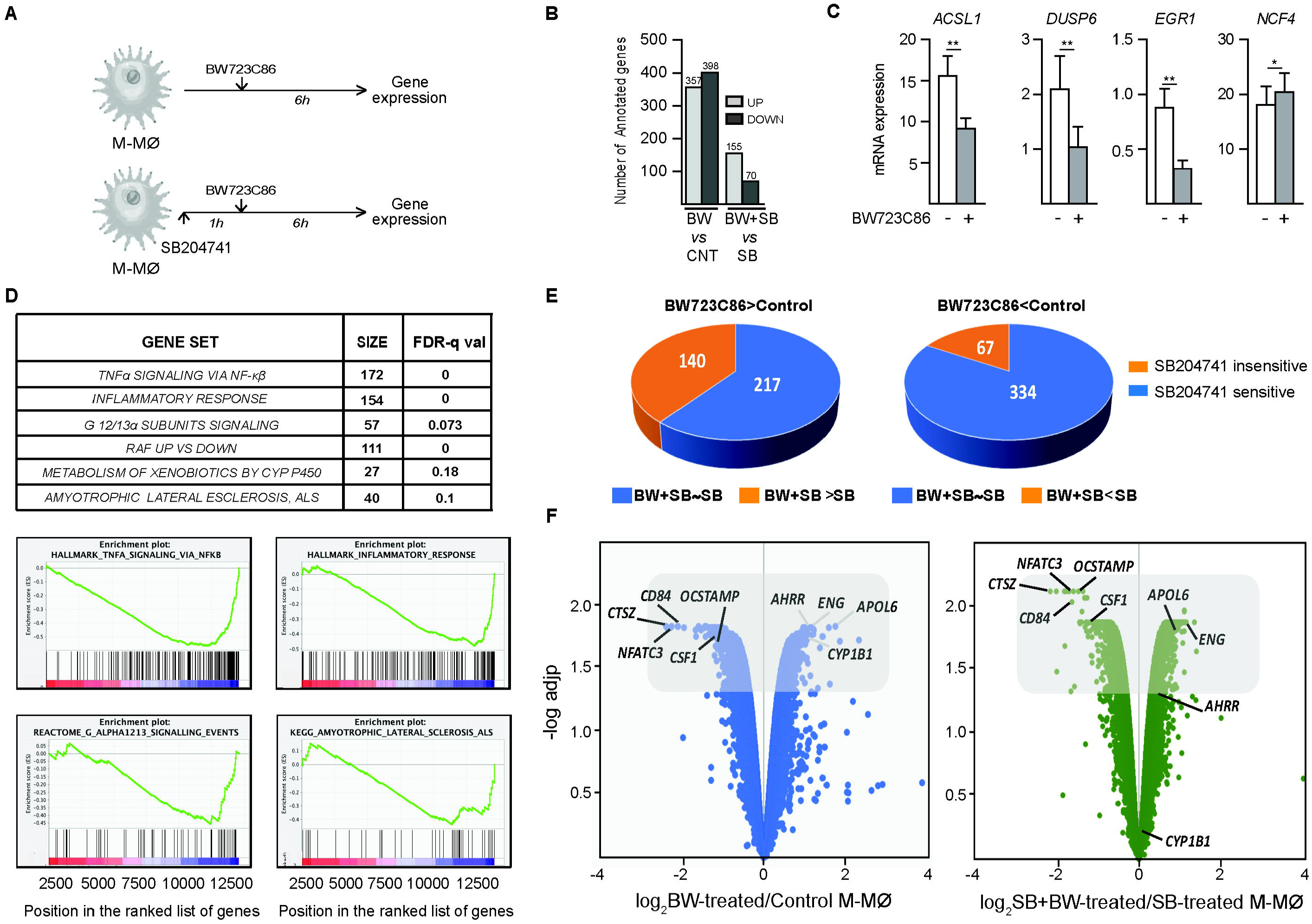
The 5-HT_2B_ agonist BW723C86 modifies the gene signature of human M-MØ. **A.** Experimental design of the gene profiling experiments. **B**. Number of annotated genes whose expression is significantly (*adj p*<0.002) upregulated (UP) or downregulated (DOWN) in M-MØ after exposure to BW723C86 (BW, 6h) in the absence (BW *vs* Control, CNT) or presence of SB204741 (SB) (BW+SB *vs* SB). **C.** Expression of the indicated genes in non-treated (−) or BW723C86-treated (6h) (+) M-MØ. Results are expressed as the mRNA level of each gene relative to the *TBP* mRNA level in the same sample. Mean and SEM of seven independent experiments is shown (*, *p* < 0.05; **, *p* < 0.01). **D.** GSEA analysis of the *t* statistic–ranked list of genes obtained from the BW723C86-treated M-MØ *versus* control M-MØ limma analysis, using the Molecular Signatures Database available at the GSEA webpage (http://software.broadinstitute.org/gsea/index.jsp) ^83^. Size and FDR *q-values* for the shown gene sets are shown at the top of the panel. **E.** Number of SB204741-sensitive and SB204741-insensitive annotated genes within the group of genes significantly (*adj p*<0.002) upregulated (BW723C86>Control, left circle) or downregulated (BW723C86<Control, right circle) by BW723C86. **F.** Volcano plot representation of the microarray data. Gene expression profiles of BW-treated M-MØ versus control M-MØ (left panel) and SB+BW-treated M-MØ versus SB-treated M-MØ (right panel) are plotted according to the log_2_ fold change (X axis) and log_10_ adjusted p-value (Y axis). In both plots, the shaded area includes the genes with *adj p*<0.05. Some of the differentially expressed genes in each case are indicated (Note the distinct relative position of *CYP1B1* and *AHRR* in both plots).

To identify genes whose expression is regulated by BW723C86 in a 5-HT_2B_-dependent manner, the transcriptional effects of the 5-HT_2B_ agonist were also determined in the presence of SB204741 (BW723C86 + SB204741) (Figure 1A, Supplementary Table IV), a selective 5-HT_2B_ antagonist with 100-fold selectivity over 5-HT_2C_ and 5-HT_2A_ ^85^ that is widely used *in vivo* ^9,61,63,90,96^. We reasoned that the genes significantly modulated by BW723C86, but not by BW723C86+SB204741, would represent *bona fide* 5-HT_2B_-regulated genes. In the presence of the antagonist SB204741, BW723C86 significantly (*adj p<0.002*) increased the expression of 155 annotated genes and downregulated the expression of 70 genes (Figure 1B, Supplementary Table V). Comparison of the BW723C86-induced transcriptional changes in the absence or presence of the antagonist (Figure 1E, F) revealed that the antagonist SB204741 prevents the BW723C86-mediated upregulation of 217 genes, and the BW723C86-mediated downregulation of 334 genes (Figure 1E, Supplementary Table VI). Therefore, 5-HT_2B_ mediates the augmented expression of 60% of the genes upregulated by BW723C86, and the diminished expression of 84% of the genes downregulated by BW723C86 (Figure 1E, Supplementary Table VI). Further, these results point towards a 5-HT_2B_-independent effect of BW723C86, as it modulates the expression of 207 genes (140 upregulated, 67 downregulated) even in the presence of the 5-HT_2B_ antagonist. As representative examples, the upregulation of *CYP1B1* and *AHRR* by BW723C86 was abrogated by SB204741, whereas SB204741 did not alter the modulation of *ENG* or *APOL6* expression by BW723C86 (Figure 1F). Altogether, these results demonstrate that BW723C86 modifies the macrophage transcriptome through 5-HT_2B_-dependent (SB204741-sensitive) and 5-HT_2B_-independent mechanisms.

### BW723C86 modifies the LPS-induced cytokine and gene profile

As GSEA revealed that BW723C86 downregulates the “TNFα Signaling via NFκβ” and “Inflammatory Response” gene sets (Figure 1D), we next assessed the ability of BW723C86 to alter macrophage responses to an inflammatory stimulus like LPS (Figure 2A). BW723C86 did not modify basal cytokine production by M-MØ (Figure 2B and not shown), while it significantly reduced TNFα production, and increased CCL2 release, from LPS-treated M-MØ (Figure 2B). A similar trend was observed with α-Methyl Serotonin (AMS), a less selective 5-HT_2B_ agonist ^77^, although its effects did not reach statistical significance (data not shown). Importantly, the effect of BW723C86 on cytokine production by LPS-activated M-MØ was blunted by the 5-HT_2B_ antagonist SB204741 (Figure 2B), thus confirming that BW723C86 modifies the LPS-induced cytokine profile via 5-HT_2B_ engagement. In fact, BW723C86 significantly impaired the acquisition of the characteristic transcriptome of LPS-stimulated M-MØ (GSE99056) ^97^, an effect that was abolished by the 5-HT_2B_ antagonist SB204741 (Figure 2C). Thus, engagement of 5-HT_2B_ by BW723C86 conditions human macrophages for altered responses to LPS.

**Figure 2.**
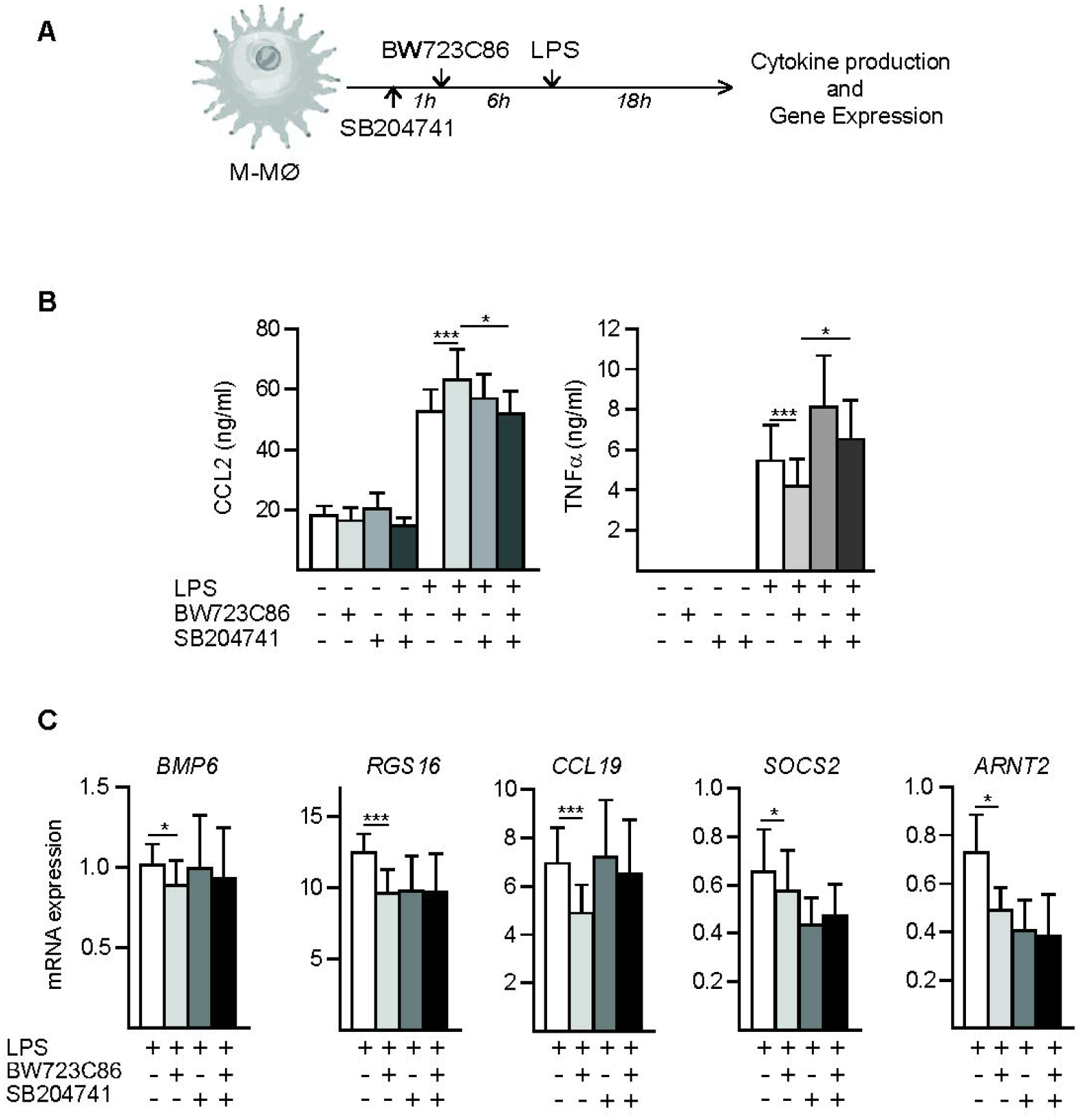
The 5-HT_2B_ agonist BW723C86 conditions human macrophages for altered LPS-stimulated cytokine and gene expression. **A.** Experimental design to assess the effect of BW723C86 and SB204741 on LPS-stimulated cytokine production and gene expression in M-MØ. **B.** Production of CCL2 and TNFα from LPS-stimulated (18h) M-MØ that had been non-treated (−) or exposed (6 h) to BW723C86, SB204741 or both (+), as determined by ELISA. Mean and SEM of 10/12 independent experiments are shown (*, *p* < 0.05; ***, *p* < 0.001). **C**. Expression of the indicated genes in LPS-stimulated (4h) M-MØ that had been non-treated (−) or exposed (6 h) to BW723C86, SB204741 or both (+), as determined by qRT-PCR. Results are expressed as the mRNA level of each gene relative to the level of *TBP* mRNA in the same sample. Mean and SEM of six independent experiments is shown (*, *p* < 0.05; ***, *p* < 0.001).

### BW723C86 upregulates the expression of AhR target genes via 5-HT_2B_

Unsupervised hierarchical clustering and ENRICHR analysis (http://amp.pharm.mssm.edu/Enrichr/) ^81,82^ on BW723C86-regulated genes (adj *p*<0.002) revealed an enrichment of the “Aryl Hydrocarbon Receptor Pathway_Homo sapiens_WP2873” and “Aryl Hydrocarbon Receptor_Homo sapiens_WP2586” gene sets (Figure 3A), suggesting that BW723C86 might influence the AhR signaling pathway. Indeed, GSEA also revealed an enrichment of the “Metabolism of xenobiotics by Cytochrome P450” gene set (Figure 3B), and of the “Ahr_UP” gene set, which contains genes upregulated by the Aryl Hydrocarbon receptor (AhR) ^98^, the major regulator of the biological responses to xenobiotics ^99^ (Figure 3B). In fact, the genes modulated by BW723C86 in an 5-HT_2B_-dependent manner included paradigmatic AhR target genes like *AHRR* and *CYP1B1* ^100^, as well as genes upregulated by the AhR agonists Benzopyrene or VAF-347 ^98,101^ and whose regulatory regions are bound by AhR (http://iregulon.aertslab.org) (Supplementary Table VI).

Based on these findings, we assessed the ability of BW723C86 to upregulate the expression of xenobiotic response genes, and found that exposure to BW723C86 significantly upregulates AhR target genes (*AHRR*, *CYP1B1*, *SERPINB2*, *TIPARP*, *EREG*) ^98,100^ (Figure 3C). Their upregulation was detected as early as 2 hours after addition of BW723C86 (Figure 3D), observed at BW723C86 concentrations between 0.1 μM and 10 μM (Figure 3E) and confirmed on 5-HT_2B_-expressing human hepatoma HepG2 cells (Figure 3F). In agreement with microarray data, upregulation of AhR-target genes by BW723C86 was reduced by the 5-HT_2B_ antagonist SB204741 (Figure 3G). Therefore, the BW723C86 agonist upregulates the expression of AhR target genes in human macrophages in a 5-HT_2B_-dependent manner.

**Figure 3.**
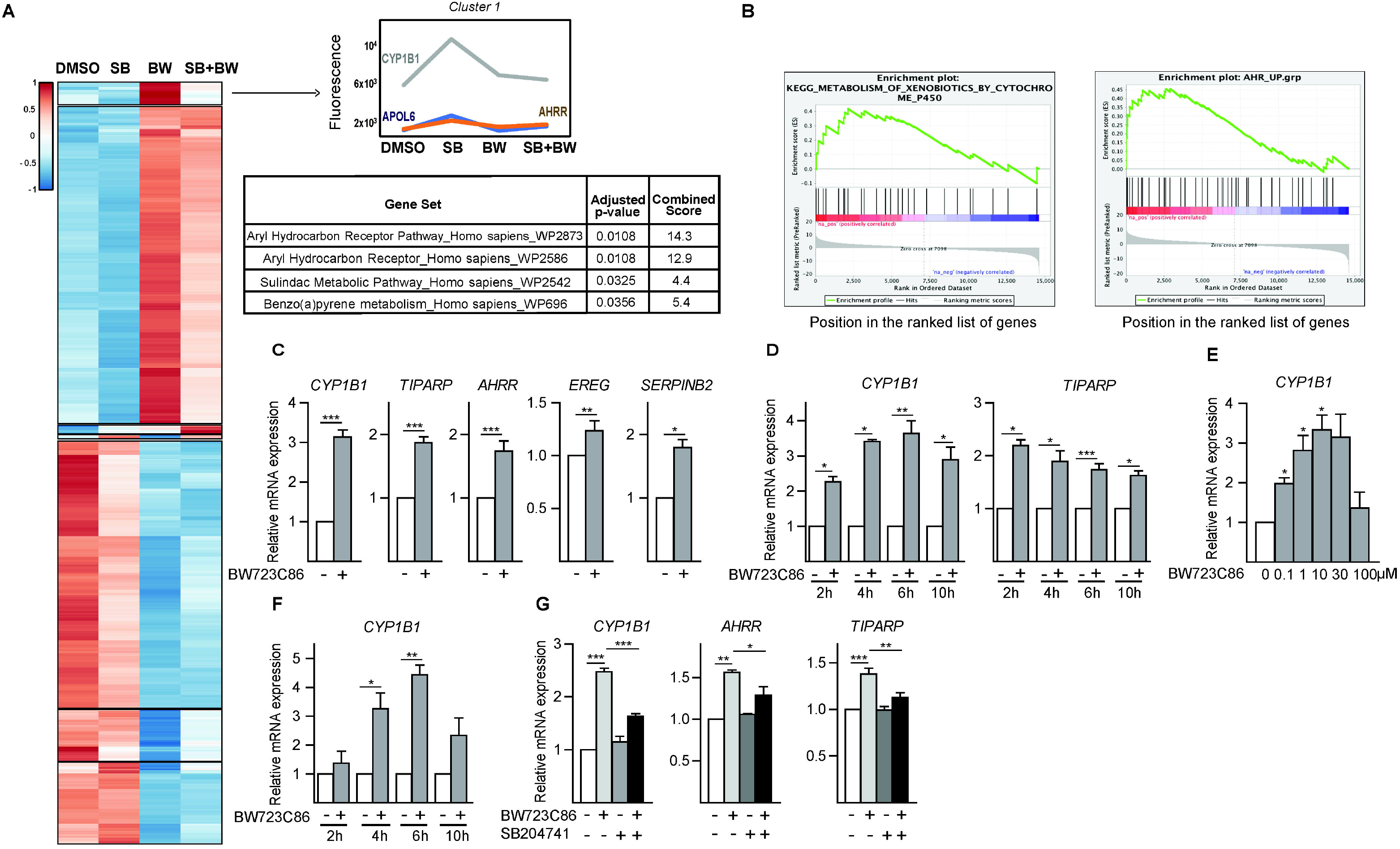
BW723C86 upregulates the expression of AhR target genes in a 5-HT_2B_ - dependent manner. **A.** Non-supervised hierarchical clustering using the ClustVis webtool (https://biit.cs.ut.ee/clustvis/) ^80^ on the mean expression level of genes significantly (adj *p*<0.002) regulated by BW723C86. For the first cluster, the pattern of expression of three representative genes and the ENRICHR data (http://amp.pharm.mssm.edu/Enrichr/) ^81,82^ on the significant enrichment of specific signaling pathway gene sets, are shown. **B.** Gene set enrichment analysis on the “t statistic-ranked” list of genes obtained from the BW723C86-treated M-MØ versus untreated-M-MØ limma analysis, using the indicated gene sets from the GSEA webpage (http://software.broadinstitute.org/gsea/index.jsp) ^83^ (upper panel) or retrieved from ^98^ (lower panel). **C.** Relative expression of the indicated genes in non-treated (−) and BW723C86-treated M-MØ (6h) (+) (n=7-11; *, *p* < 0.05; **, *p* < 0.01; ***, *p* < 0.001). **D.** Relative expression of *CYP1B1* and *TIPARP* in non-treated (−) and M-MØ treated with BW723C86 (+) for the indicated periods of time (n=3; *, *p* < 0.05; **, *p* < 0.01; ***, *p* < 0.001). **E.** Relative expression of *CYP1B1* in non-treated (0) or M-MØ treated for 6h with the indicated concentrations of BW723C86 (n=3; *, *p* < 0.05). **F.** Relative expression of *CYP1B1* in non-treated (−) or HepG2 hepatoma cells treated with BW723C86 (+) for the indicated periods of time (n=3 *, *p* < 0.05; **, *p* < 0.01). **G**. Relative expression of *CYP1B1*, *AHRR* and *TIPARP* in non-treated (−) or M-MØ treated for 6h with BW723C86, SB204741 or both (+) (n=5; *, *p* < 0.05; **, *p* < 0.01; ***, *p* < 0.001). (**D-G)** Results are shown as the expression of each gene after the different treatments and relative to its expression in control (untreated) samples. In all cases, mean and SEM are shown.

### AhR mediates the BW723C86-induced upregulation of AhR target genes in human macrophages

To evaluate whether BW723C86 leads to AhR activation, its transcriptional effects were determined after pharmacological or siRNA-mediated inhibition of AhR. The AhR antagonist CH223191 ^102^ prevented the BW273C86-induced upregulation of both *CYP1B1* and *TIPARP*, whose expression was increased by the AhR agonist FICZ ^103^ (Figure 4A). Besides, siRNA-mediated knockdown of AhR (Figure 4B,C) completely prevented the BW273C86-induced *CYP1B1* and *TIPARP* upregulation (Figure 4D). In agreement with the role of AhR in the BW723C86-induced upregulation of AhR-target genes, reporter gene experiments revealed that BW723C86 specifically enhances the transcriptional activity of AhR in M-MØ (Figure 4E), an effect that was abrogated by the AhR antagonist CH223191 (Figure 4E). Similar results were also obtained in the 5-HT_2B_-expressing HepG2 cell line, where the BW723C86-induced AhR transcriptional activity was prevented by the 5-HT_2B_ antagonist SB204741 (Figure 4F) or the AhR antagonist CH223191 (Figure 4G). Therefore, engagement of the 5-HT_2B_ serotonin receptor by BW723C86 in human macrophages enhances the transcriptional activity of AhR and results in upregulated expression of AhR-target genes.

**Figure 4.**
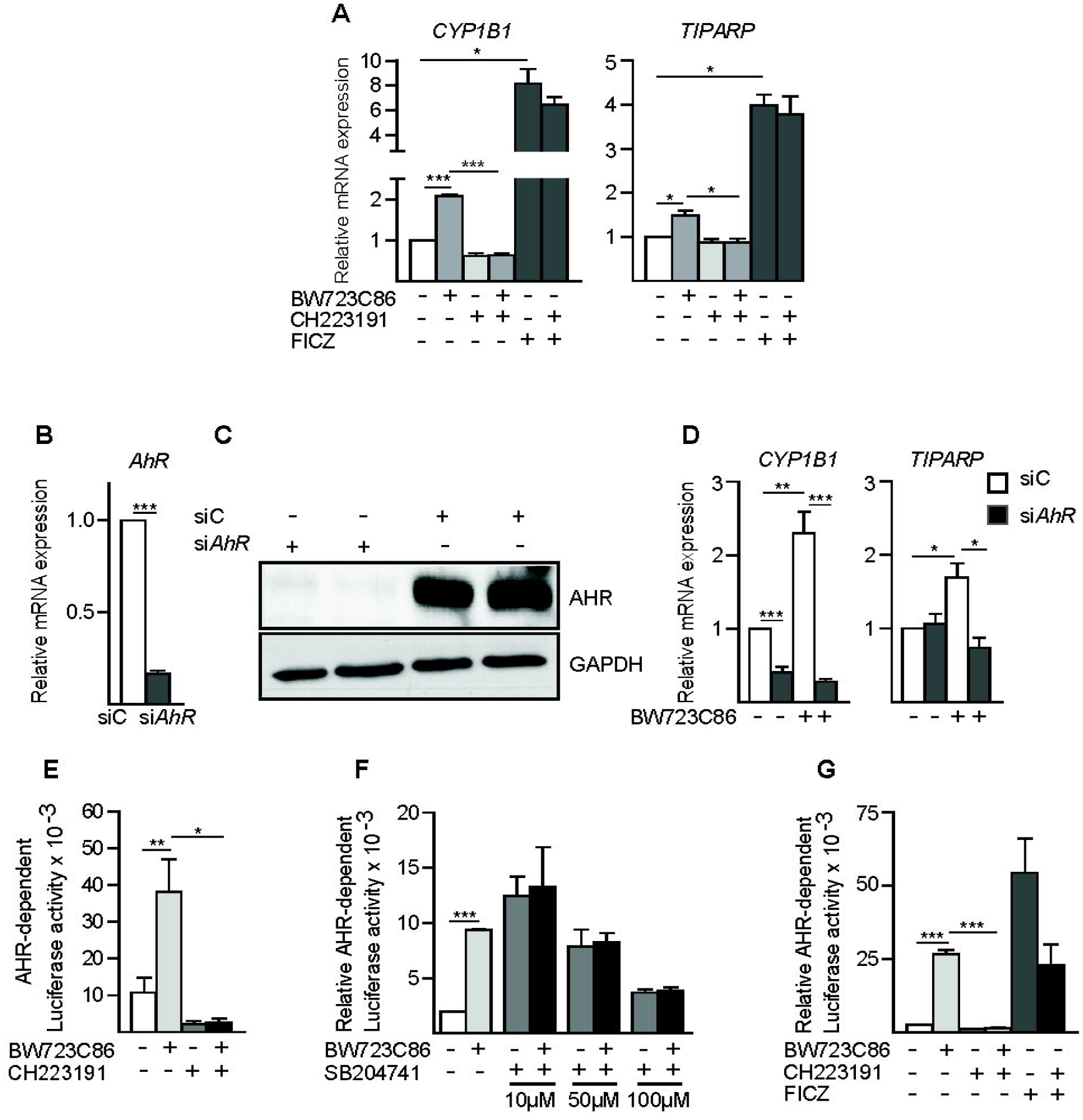
CYP1B1 upregulation by BW723C86 is dependent on 5-HT_2B_ and requires AhR activation. **A.** Relative expression of *CYP1B1* and *TIPARP* in M-MØ either untreated (−) or treated for 6h with BW723C86 (BW) in the absence or presence (+) of the AhR antagonist CH223191. M-MØ were treated in parallel with the AhR agonist FICZ (6h) for control purposes. Results are shown as the expression of each gene after the different treatments and relative to its expression in control (untreated) samples. In all cases, mean and SEM are shown (n=4; *, *p* < 0.05; ***, *p* < 0.0005). **B-C.** *AHR* mRNA (**B**) and AhR protein level (**C**) in M-MØ transfected with control siRNA (siC) or an AhR-specific siRNA (siAhR). 18 hours after transfection, M-MØ were washed and lysed for qRT-PCR (**B**) or Western blot (**C**). Six independent experiments were performed for *AHR* mRNA determination by qRT-PCR (**B**) (n=6; ***, *p* < 0.001), and Western blots from two independent knockdown experiments are shown in (**C**). **D**. Relative expression of *CYP1B1* and *TIPARP* in M-MØ transfected with control siRNA (siC) or AhR-specific siRNA (siAhR) and either left untreated (−) or treated for 6h with BW723C86 (+) (n=6; *, *p* < 0.05; **, *p* < 0.01; ***, *p* < 0.001). In **A-D**, results are shown as the expression of each gene after the different treatments and relative to its expression in control (untreated) samples. In all cases, mean and SEM are shown. **E.** AhR-dependent transcriptional activity in XRE-Luc transfected M-MØ left untreated (−) or treated with BW723C86 (14h) in the absence or in the presence of the AhR antagonist CH223191, added 1 hour before agonist addition. Mean and SEM of the AhR-dependent luciferase activity of five independent experiments is shown (n=5; *, *p* < 0.05; **, *p* < 0.01). **F.** AhR-dependent transcriptional activity in XRE-Luc-transfected HepG2 cells left untreated (−) or treated with BW723C86 (16h) in the absence or in the presence of the indicated concentrations of the 5-HT_2B_ antagonist SB204741 (n=4; ***, *p* < 0.001). **G**. AhR-dependent transcriptional activity in XRE-Luc-transfected HepG2 cells left untreated (−) or treated with BW723C86 (16h) in the absence or presence of the AhR antagonist CH223191, which was added 1 hour before agonist addition. HepG2 cells were treated with FICZ for control purposes (n=4; ***, *p* < 0.001). In **F-G**, results are shown as the mean and SEM of the Relative AhR-dependent luciferase activity of each sample (referred to the luciferase activity measured in untreated samples).

### 5-HT and other 5-HT_2B_ ligands influence the expression of BW723C86-regulated genes

To further support the involvement of 5-HT_2B_ in the ability of BW723C86 to promote AhR activation, the influence of 5-HT and other 5-HT_2B_ agonists on the expression of AhR target genes was determined. Albeit to a lower extent than BW723C86, the potent 5-HT_2B_ agonist 6-APB (10nM-1μM) upregulated *CYP1B1* expression in M-MØ (Figure 5A). Similarly, the 5-HT_2B_ agonist alpha-Methyl-Serotonin (AMS) enhanced macrophage *CYP1B1* expression in a dose-dependent manner, and its effect was abolished by the SB204741 antagonist (Figure 5B). Therefore, 5-HT_2B_ agonists upregulate AhR target gene expression in a 5-HT_2B_-dependent manner in human macrophages. Next, we tested whether 5-HT has an impact on the BW723C86-induced transcriptional effects. As expected ^39^, 5-HT treatment upregulated the expression of *PDE2A* and *TREM1* but did not modify the expression of *CYP1B1* (Figure 5C). However, the BW723C86-mediated upregulation of *CYP1B1* was prevented in the presence of 100 μM 5-HT that, by contrast, did not affect the ability of BW723C86 to downregulate *OCTSAMP* expression (Figure 5C). Therefore, 5-HT impairs the ability of the 5-HT_2B_ agonist BW723C86 to activate the AhR transcriptional activity in human macrophages.

**Figure 5.**
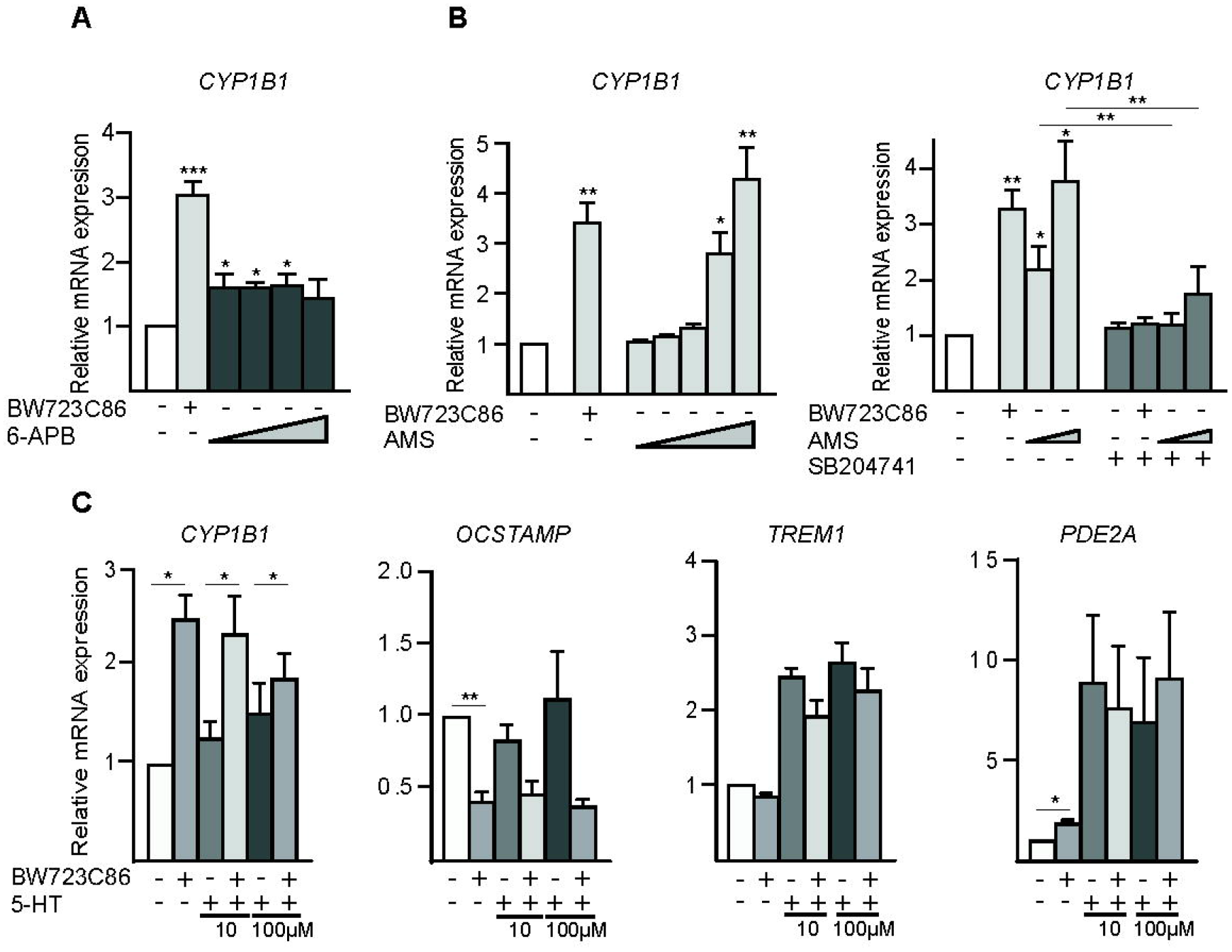
AhR activation in human macrophages is also triggered by other 5-HT_2B_ ligands and is prevented by serotonin. **A.** Relative expression of *CYP1B1* in M-MØ treated for 6h with DMSO (vehicle, -), BW723C86 (+) or with increasing concentrations of APB (10 nM, 50 nM, 100 nM or 1 μM) (n=9). **B.** Relative expression of *CYP1B1* in non-treated (−) or M-MØ treated for 6h with BW723C86 (+) or with increasing concentrations of α-Methyl 5HT (AMS) (10 nM, 20 nM, 50 nM,100μM or 500 μM) (left panel) (n=6), and either in the absence or presence of SB204741 using 100μM or 500 μM AMS (right panel) (n=4). (**A-B)** Results are shown as the expression of each gene after the different treatments and relative to its expression in control (untreated) samples, and mean and SEM are shown (*p* < 0.05; **, *p* < 0.01; ***, *p* < 0.001). **C**. Relative expression of the indicated genes in M-MØ either untreated (−) or treated for 6h with 10 μM BW723C86, 5-HT (10 μM or 100 μM) or both (n=4). Shown is the expression of each gene after the different treatments and relative to its expression in control (untreated) samples. Mean and SEM are shown (*p* < 0.05; **, *p* < 0.01; ***, *p* < 0.001).

### 5-HT2B-independent effects of BW723C86

The identification of 207 genes whose regulation by BW723C86 is not affected by the 5-HT_2B_ antagonist SB204741 (Figure 1E,F) suggested the existence of 5-HT_2B_-independent effects of BW723C86. As an example, the BW723C86-mediated downregulation of *OCSTAMP* was not altered by 5-HT or the 5-HT_2B_ antagonist. Along the same line, while the BW723C86-mediated changes in *CYP1B1* and *CSF1* expression was inhibited by the AhR antagonist CH223191, AhR inhibition had no influence of the downregulation of *EGR1* and *OCSTAMP* by BW723C86 (Figure 6). Altogether, these results confirmed the existence of transcriptional changes induced by BW723C86 that are AhR-independent and not prevented by the 5-HT_2B_ antagonist SB204741 (HT_2B_-independent).

**Figure 6.**
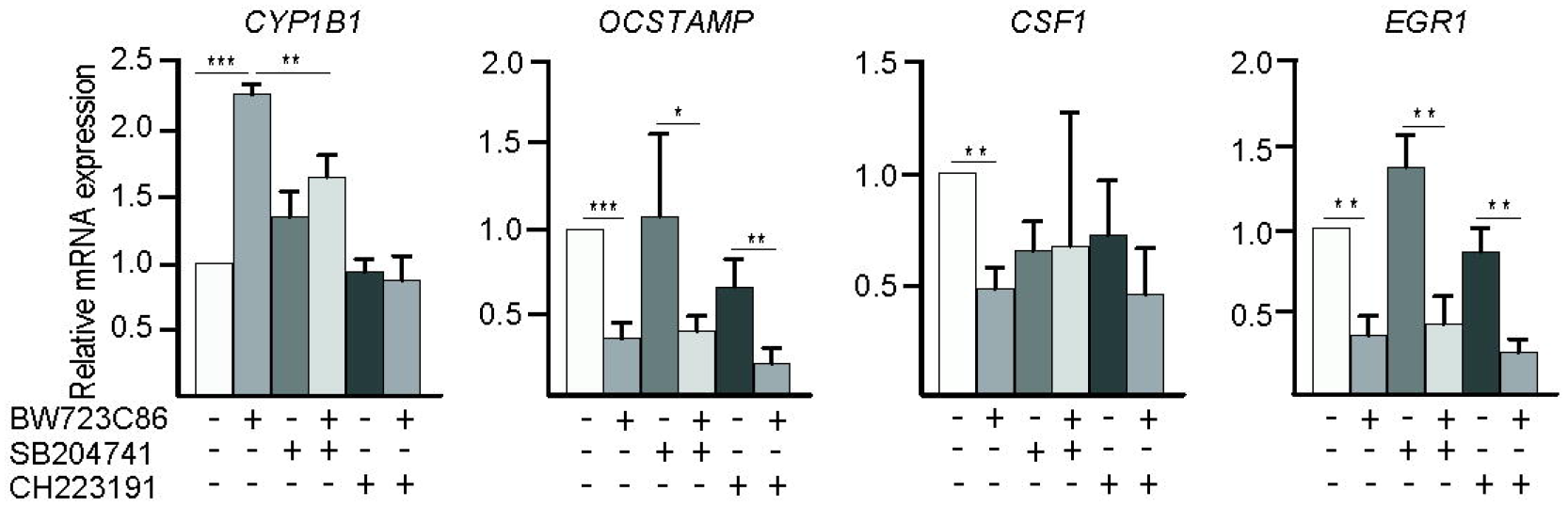
BW723C86 also regulates the macrophage transcriptome in a 5-HT_2B_-independent manner. Relative expression of the indicated genes in non-treated (−) or M-MØ treated for 6h with BW723C86 after a previous treatment (1h) with either SB204741 or CH223191 (n=5; ***, *p* < 0.001). Results are shown as the expression of each gene after the different treatments and relative to its expression in control (untreated) samples. In all cases, mean and SEM are shown.

### BW723C86 blocks osteoclast differentiation

The ability of BW723C86 to downregulate the expression of *CSF1* (which encodes M-CSF), *EGR1* (which regulates osteoclastogenesis) ^104,105^ and *OCSTAMP* (which drives osteoclast precursor fusion) ^106^ (Figure 6), together with the under-expression of the “GO Regulation of Osteoclast Differentiation” gene set in the BW723C86-M-MØ transcriptome (Supplementary Figure 1A), led us to test whether BW723C86 negatively affects the monocyte-to-osteoclast differentiation. The continuous presence of the agonist during M-CSF+RANKL-induced osteoclast differentiation significantly impaired the expression of *OCSTAMP* and *DCSTAMP*, which mediate osteoclast cell-cell fusion ^107,108^, as well as the expression of hallmark osteoclast markers like *CTSK*, *CALCR*, *CA2* and *TNFRSF11A* (which encodes the RANKL receptor) (Figure 7A). Therefore, BW723C86 inhibits the expression of genes involved in osteoclast differentiation, cellular fusion and bone resorption, a previously unnoticed action of this 5-HT_2B_ agonist.

**Figure 7.**
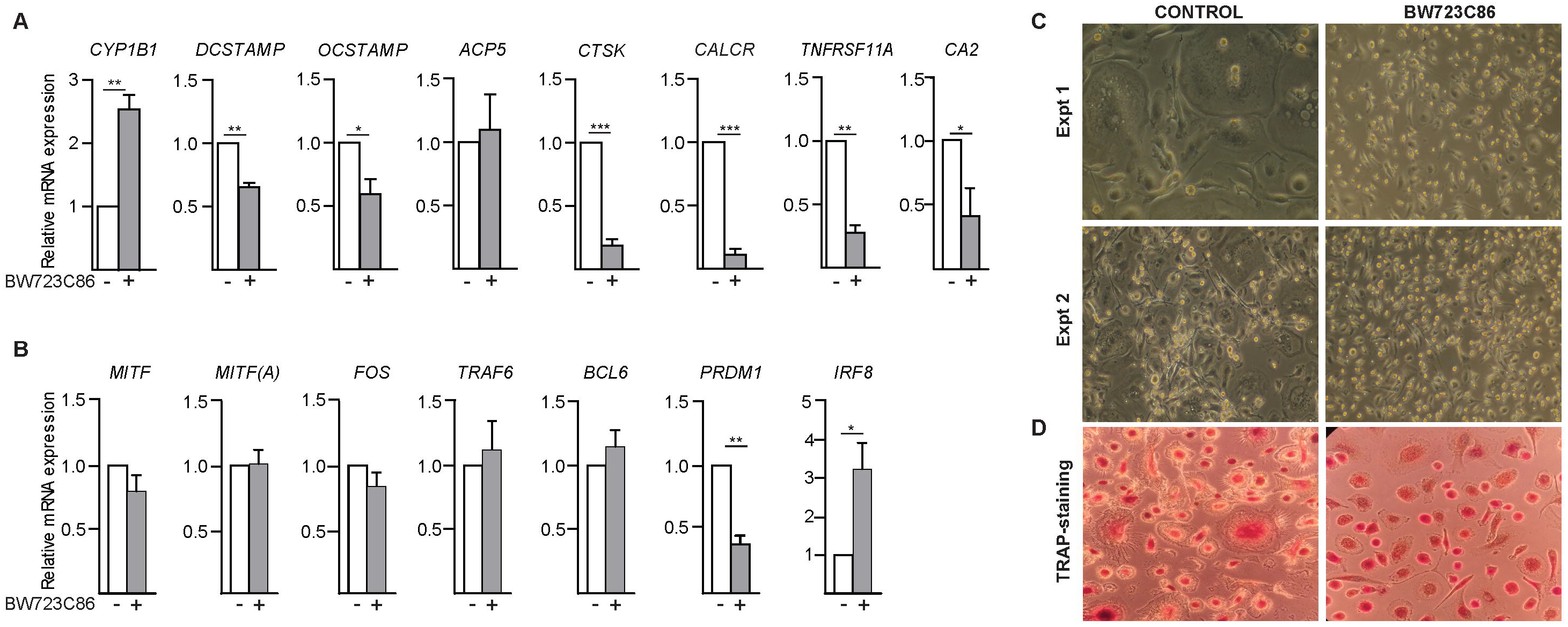
BW723C86 prevents the generation of osteoclasts from human peripheral blood monocytes. **A-B.** Expression of the indicated genes in monocyte-derived osteoclasts generated after a 12 day-culture with M-CSF and RANKL in the absence (−) or presence of BW723C86 (+) (n=4; *, *p* < 0.05; **, *p* < 0.01; ***, *p* < 0.001). Results are shown as the expression of each gene in BW723C86-treated cells and relative to its expression in control (untreated) samples. In all cases, mean and SEM are shown. **C-D.** Phase-contrast microscopy (**C**) and TRAP-staining (**D**) of untreated (CONTROL, left panels) or BW723C86-treated monocytes cultured for 12 days in the presence of M-CSF and RANKL for differentiation into osteoclasts (right panels). Two independent experiments are shown in panel **C**.

To get further insight into the inhibitory action of BW723C86 on osteoclast differentiation, we next assessed whether the agonist modulates the expression of genes coding for factors driving osteoclastogenesis. The presence of BW723C86 along osteoclastogenesis did not modify the expression of *TRAF6*, *MITF*, *FOS* and *BCL6* (Figure 7B). However, BW723C86 significantly augmented the expression of *IRF8*, whose protein product negatively regulates osteoclast differentiation ^107^, and simultaneously provoked a significant reduction in the expression of *PRDM1*, whose encoded protein (Blimp1) represses the expression of negative regulators of osteoclast differentiation ^107,109,110^ (Figure 7B). In fact, exposure of M-MØ to BW723C86 for 6 hours sufficed to significantly downregulate the expression of *PRDM1* (Supplementary Table II). In consonance with these results, the continuous presence of BW723C86 greatly impaired the generation of “ruffled border”-rich multinucleated cells during the M-CSF+RANKL-driven osteoclast differentiation, as seen by light microscopy (Figure 7C) and after TRAP staining (Figure 7D). Therefore, BW723C86 has an inhibitory effect on the ability of human monocytes to differentiate into osteoclasts at the transcriptional and morphological level, an effect that correlates with its ability to down-regulate *PRDM1* expression and to enhance the expression of *IRF8*.

## DISCUSSION

The polarization-specific expression of 5-HT_2B_, an unintended target of commonly used drugs, prompted us to dissect the transcriptional consequences of 5-HT_2B_ activation in human macrophages through the use of the 5-HT_2B_ agonist BW723C86, which exhibits good selectivity for 5-HT_2B_ ^85^, and the 5-HT_2B_ antagonist SB204741, with exerts selectivity over 5-HT_2C_ and 5-HT_2A_ ^85^. We now provide evidences that 5-HT_2B_ engagement shapes the macrophage transcriptome partly via activation of AhR ^99^ and that BW723C86 impairs the osteoclastogenic differentiation of monocytes at the morphological and transcriptional level, thus emphasizing the role of 5-HT_2B_ in myeloid cell differentiation. The 5-HT_2B_-AhR axis extends the range of signaling pathways initiated upon 5-HT receptor engagement and constitutes a novel point of convergence between endogenous and exogenous agents with capacity to modulate inflammatory responses.

The link between 5-HT_2B_ and AhR activation is compatible with 5-HT_2B_ favouring an anti-inflammatory polarization because AhR enhances IL-10 expression and limits pro-inflammatory cytokine expression in macrophages ^111^. The transcriptional profile promoted by BW723C86 in human macrophages fits with the known signaling ability of 5-HT_2B_ ^112^, as it shows a significant enrichment of genes involved in G protein signaling (Figure 1D). Like 5-HT, BW723C86 has been shown to induce hepatocyte proliferation via PLC activation, EGFR phosphorylation and subsequent mTOR, PI3K, ERK and PLA2 activation ^113^, and these signaling molecules have been recurrently linked to 5-HT_2B_ activation ^112^. By contrast, the unexpected link of 5-HT_2B_ to AhR activation constitutes an addition to the plethora of cell-specific intracellular signaling pathways activated by 5-HT_2B_, and its significance is emphasized by the finding that the 5-HT_2B_-AhR link is also functional in the 5-HT_2B_-expressing HepG2 hepatoma cell line. As a whole, our results point to AhR and AhR-target genes (e.g., *CYP1B1*, *AHRR*, *TIPARP*, …) as effectors of the 5-HT_2B_ receptor in human macrophages.

The transcriptional effects of BW723C86 on human macrophages differ considerably from those triggered by 5-HT, most of which can be mimicked or abrogated by 5-HT_7_ agonists or antagonists, respectively ^39^. Thus, although 5-HT_2B_ and 5-HT_7_ are preferentially expressed by anti-inflammatory human macrophages ^40^ and promote a pro-fibrotic phenotype ^19,39,47,114,115^, their individual engagement leads to different transcriptional outcomes, with AhR-activating ability being restricted to 5-HT_2B_. The failure of 5-HT to activate AhR in human macrophages might rely on the fact that 5-HT can bind to both 5-HT_2B_ and 5-HT_7_, and can therefore initiate intracellular signaling from both receptors. Ligation of 5-HT_7_ by 5-HT triggers protein kinase A (PKA)-dependent signaling and transcription in macrophages ^39^ and other cell types ^116^, and results in the expression of PKA-regulated genes like *PDE2A* ^39^. Since PKA activation (or increased cAMP levels) represses AhR-dependent gene expression ^117,118^, it is therefore possible that the binding of 5-HT to 5-HT_7_, and the resulting PKA activation ^39,116^, might prevent the activation of AhR secondary to 5-HT binding to 5-HT_2B_. In line with this hypothesis, it is worth noting that *PDE2A* gene expression in macrophages is greatly upregulated after 5-HT binding to 5-HT_7_ ^39^, and that PDE2A interaction with the immunophilin-like protein XAP2 (a member of the molecular complex that retains unliganded AhR in the cytoplasm) inhibits dioxin-induced AhR nuclear translocation and transcription in hepatocytes ^119^.

A recent report has illustrated that, in intestinal epithelial cells, 5-HT activates AhR and the expression of AhR target genes in a manner that is independent on 5-HT receptors but mediated by the serotonin transporter (SERT) ^120^. The BW723C86-mediated activation of AhR in macrophages (here reported) is unrelated to the 5-HT/SERT/AhR link because: 1) it can be inhibited by an antagonist of 5-HT_2B_; 2) M-MØ are devoid of SERT expression ^84^; and 3) 5-HT does not upregulate *CYP1B1* expression in human monocyte-derived macrophages. However, the existence of these two pathways (5-HT/SERT/AhR in intestinal epithelial cells and BW723C86/5-HT_2B_/AhR in macrophages) fits with the AhR-activating ability of tryptophan catabolites ^121^ and supports the importance of 5-HT and related molecules in modulating AhR activity and, hence, inflammatory responses. In this regard, and besides a receptor for environmental toxins ^99^, AhR mediates vascular system development, immune system polarization and resolution of inflammatory responses ^122,123^, processes where 5-HT_2B_ has been also implicated ^16,70,71,93^. Thus, since 5-HT_2B_ agonists (and specially BW723C86) have been widely used *in vivo* to support the participation of 5-HT_2B_ in several animal models of disease ^73,74,86,87,89-91,124-126^, our results suggests the possibility that AhR might have influenced some of these previous results and might have even contributed to the pathological effects of deregulated 5-HT_2B_ expression or function. As an example, since AhR mediates IL-10 production and promotes immunological tolerance after macrophage capture of apoptotic cells ^127^, AhR might contribute to the anti-inflammatory effect of 5-HT_2B_ in mouse microglia ^71^ and human monocytes ^72^. On a related note, the assessment of the extent of the transcriptional action of BW723C86 evidenced that the effects of the agonist on the expression of more than 200 genes (including some encoding regulators of osteoclastogenesis) are not prevented by the 5-HT_2B_ antagonist SB204741 (Figure 1). This finding indicates the existence of 5-HT_2B_-independent effects of BW723C86 and, in line with the above comments, suggest that some of the actions previously assigned to BW723C86 *in vivo* and *in vitro* might be actually unrelated to (independent on) the ability of BW723C86 to interact with 5-HT_2B._

## Supporting information

Supplementary Table III

## ACKNOWLEDGEMENTS

This work was supported by grants from Ministerio de Economía y Competitividad (SAF2014-52423-R and SAF2017-83785-R) to MAV and ALC, Grant 201619.31 from Fundación La Marató/TV3 to ALC, and Red de Investigación en Enfermedades Reumáticas (RIER, RD16/0012/0007), and cofinanced by the European Regional Development Fund “A way to achieve Europe” (ERDF). MCE and IR were funded by FPI predoctoral fellowship (BES-2009-021465 and BES-2015-071337, respectively) from Ministerio de Economía e Innovación.

## AUTHORSHIP CONTRIBUTIONS

CN, IR, MCE, EI and BA performed research and analyzed data; CN, IR, MCE, MAV and ALC designed the research and analyzed data; ALC wrote the paper.

---

***Supplementary Figure 1.* A.** Selected GSEA on the “t statistic-ranked” list of genes obtained from the BW723C86-treated M-MØ versus untreated-M-MØ limma analysis, using the C5 collection within the Molecular Signatures Database (MSigDB) gene sets (http://software.broadinstitute.org/gsea/index.jsp). **B.** GSEA on the “t statistic-ranked” list of genes obtained from the BW723C86-treated M-MØ versus untreated-M-MØ limma analysis, using the “Proinflammatory gene set” (left) and “Anti-inflammatory gene set” (right) previously defined ^84,95,128^.

