## Supplementary figures and images for "5-HT_2B_ serotonin receptor agonist BW723C86 shapes the macrophage gene profile via AhR and impairs monocyte-to-osteoclast differentiation"

### Supplementary Figure 1

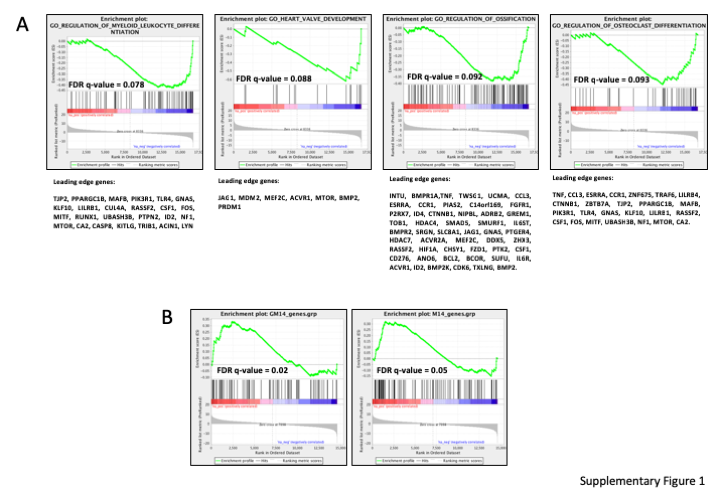
